# Dynamic microtubules drive yolk-cytoplasm segregation in the syncytial *Drosophila* embryo

**DOI:** 10.64898/2026.09.02.748760

**Authors:** Sameer Thukral, Pampa Dey, Bivash Kaity, S Sanjana, Shashwat Goswami, Yu-Chiun Wang, Mithun K. Mitra, Amitabha Nandi, Richa Rikhy

**Affiliations:** Indian Institute of Science Education and Research, Pune, India; Laboratory for Epithelial Morphogenesis, RIKEN-BDR, Kobe, Japan; Department of Physics, Indian Institute of Technology Bombay, Mumbai, India

**Keywords:** Yolk-cytoplasm segregation, microtubules, syncytial blastoderm, *Drosophila* embryogenesis, cytoskeletal dynamics

## Abstract

Yolk-cytoplasm segregation is among the earliest spatial organization events in the developing embryo of many oviparous animals. The segregation process is intimately linked to early embryonic cleavage and pattern formation, and exhibits a wide range of spatial and temporal diversity. However, the underlying cytoskeletal mechanism remains largely unknown, except for a small number of species. Using quantitative live imaging, we investigated yolk segregation in the *Drosophila* embryo during the syncytial nuclear cycles 11–14. We find that the yolk vesicles move progressively inward in spatial and temporal coordination with the inward expanding microtubule networks that are nucleated from centrosomes positioned at the cortex, whereas cortical actin meshwork remains spatially restricted. Using the *gnu* RNAi embryo to decouple nuclear migration and division from cytoskeletal dynamics, we establish causality with targeted pharmacological disruption and find that microtubule dynamics is required for yolk segregation, while depolymerization of actin has no discernible effect. In support of a mechanism of growth-propelled passive displacement, microtubule plus end comets come in apparent contact with yolk vesicles, and injected, inert microbeads are displaced towards the embryo center presumably by the same pushing force. These findings identify microtubule polymerization as a predominant driver of yolk-cytoplasm segregation in *Drosophila* and suggest that diverse cytoskeletal mechanisms evolved to accomplish this crucial reorganization process.

## Introduction

Egg-laying, or oviparous animals maternally assemble and deposit a nutrient store in the developing oocyte. This store, called the yolk, supplies energy and molecular building blocks, thereby sustaining embryonic development until the embryo hatches and acquires the capability to self-feed ^1^. In the early embryo, the yolk undergoes a partitioning process – nearly invariably in all oviparous animals – to become separated from the rest of the cytoplasm. This process of yolk-cytoplasm segregation marks one of the earliest organizational events in the developing embryo, separating the embryo into the blastoderm and the yolk^1^. Past studies in a number of different organisms have suggested that defective yolk-cytoplasmic organization could have developmental consequences during late stage embryonic development ^2–4^, making it a critical determinant of embryonic success. Additionally, yolk-cytoplasm segregation has emerged as an important model for the study of intracellular compartmentalization and maintenance of cytoplasmic inhomogeneities, topics with critical implications on the physical principles of self organization in living systems ^5^.

Yolk-cytoplasm segregation is spatially and temporally intertwined with early embryonic processes that include positioning and mobility of the spindles ^6^, timing and patterning of early embryonic cleavage cycles ^7^, mobility and polarization of maternally-derived cell fate and patterning determinants ^8^, and the morphogenesis of gastrulation ^9,10^. Given such an intimate connection, it has been hypothesized that yolk segregation influences or acts as an evolutionary constraint on the modes of early embryonic development ^9,11,12,13^. Despite that, the underlying process has only been studied mechanistically in a small number of species. Notably, in the zebrafish embryo, one of the best characterized systems thus far ^7^, the cell cycle machinery that drives the first embryonic cleavage initiates a wave of bulk actin polymerization.

This wave sweeps across the oocyte, pulling the cytoplasm (ooplasm) toward the animal pole, while pushing yolk granules down toward the vegetal pole by forming actin comets on the surface of yolk granules ^7^. Beyond zebrafish, remodeling and rearrangement of the cytoskeleton, or large-scale cytoplasmic streaming, has been found to be associated with yolk-cytoplasm segregation in other organisms, such as the nematodes ^14,6^, ascidians^15^, and amphibians^16,17^. The spatial features, modes and timing of segregation, as well as how it is linked to the early embryonic developmental timetable, vary widely within a phylogenetic clade ^15^ and across species. And yet, it remains unclear whether mechanisms that drive the segregation of yolk from cytoplasm are diverse or conserved.

The *Drosophila* embryo develops as a syncytium, where the early embryonic nuclei divide 13 times in a shared cytoplasm that contains three major yolk components – the protein rich yolk vesicles, lipid droplets and glycogen granules ^4^. Contrasting with the zebrafish embryo, where yolk segregation takes place prior to the first embryo cleavage, the initial nuclear division cycles (NCs) occur deep within the syncytial embryo, in the absence of any spatial separation or compartmentalization of the yolk, syncytial nuclei and mitotic apparatus. From NC9 onward, the majority of embryonic nuclei begin to migrate to the embryo cortex. This migration coincides with the formation of an optically clear cortical cytoplasm that widens progressively, as yolk components, especially the yolk vesicles, partition into an optically opaque mass at the center of the embryo^18^. Previous work noted a correlation between the movements of the yolk and nuclei ^18^. The underlying mechanics of the yolk movements, and in particular the cytoskeletal forces that drive these dynamics, however, remain largely unknown.

Here we use quantitative live imaging and targeted cytoskeletal perturbations to characterize the dynamics of yolk-cytoplasm segregation in *Drosophila*. We find that the yolk vesicles move progressively inward during NC11-14. In the *giant nuclei* (*gnu*) RNAi embryos where the cytoskeletal networks are established at the embryo cortex despite defective nuclear migration, pharmacological perturbations reveal that microtubules (MTs) play a predominant role in yolk-cytoplasm segregation, while actin depolymerization has no detectable effect. Our work establishes a novel and alternative mechanistic principle for yolk-cytoplasm segregation, enriching our appreciation of the evolutionary diversity of cytoplasmic organization during early embryonic development.

## Results

### The yolk vesicles progressively move towards the embryo center

To characterize yolk-cytoplasm segregation in the early *Drosophila* embryo, we imaged the dynamics of the yolk vesicles, a principal component of the yolk, and quantified their size, dynamics, position and density. Two complementary approaches were employed to visualize the yolk vesicles: 1) a negative labeling approach using a maternally deposited cytoplasmic mRFP, revealing the yolk vesicles as dark circular objects (Figure 1A, 1B, Movie S1); 2) a direct labeling approach using GFP-tagged Yolk protein 1 (Yp1-GFP), which fills these vesicles, providing a validation to results obtained with the negative labeling method (Figure S1A, S1B, Movie S2; see Figure S1C). The size of the vesicles is estimated to range from 1 to 9 μm in diameter, with a median of 3.4 μm (Figure S1D). The vesicles progressively move toward the center of the embryo from NC11 to NC14 (Figure 1B). Kymograph based analysis revealed that the yolk-cytoplasm boundary moves at a speed of 0.3 **±** 0.1 μm/min away from the embryo cortex over these last four NCs (Figure 1C, 1D, S1E-G, “KM”). To cross validate these results, we employed two additional methods: First, the vesicles were imaged *en face* in 3D, segmented to measure yolk occupancy across a z-plane, followed by fitting to a mass conservation equation to calculate the flux of the vesicles (Figure 1E, 1F, S1H-I, “MCE”; see Methods for details); Second, the vesicles were imaged in 2D, segmented and tracked using the FIJI tracking suite, TrackMate ^19^ (Figure 1F, S1F, “TM”). These two methods yielded comparable vesicle velocities (0.4 **±** 0.02 and 0.4 **±** 0.09 μm/min, respectively; Figure 1F, S1F). Quantitative analyses revealed a steady increase in the distance of the vesicles from the embryo surface (Figure 1G) and an increase in the density of the vesicles in the embryo center across these four NCs (Figure 1H).

**Figure 1.**
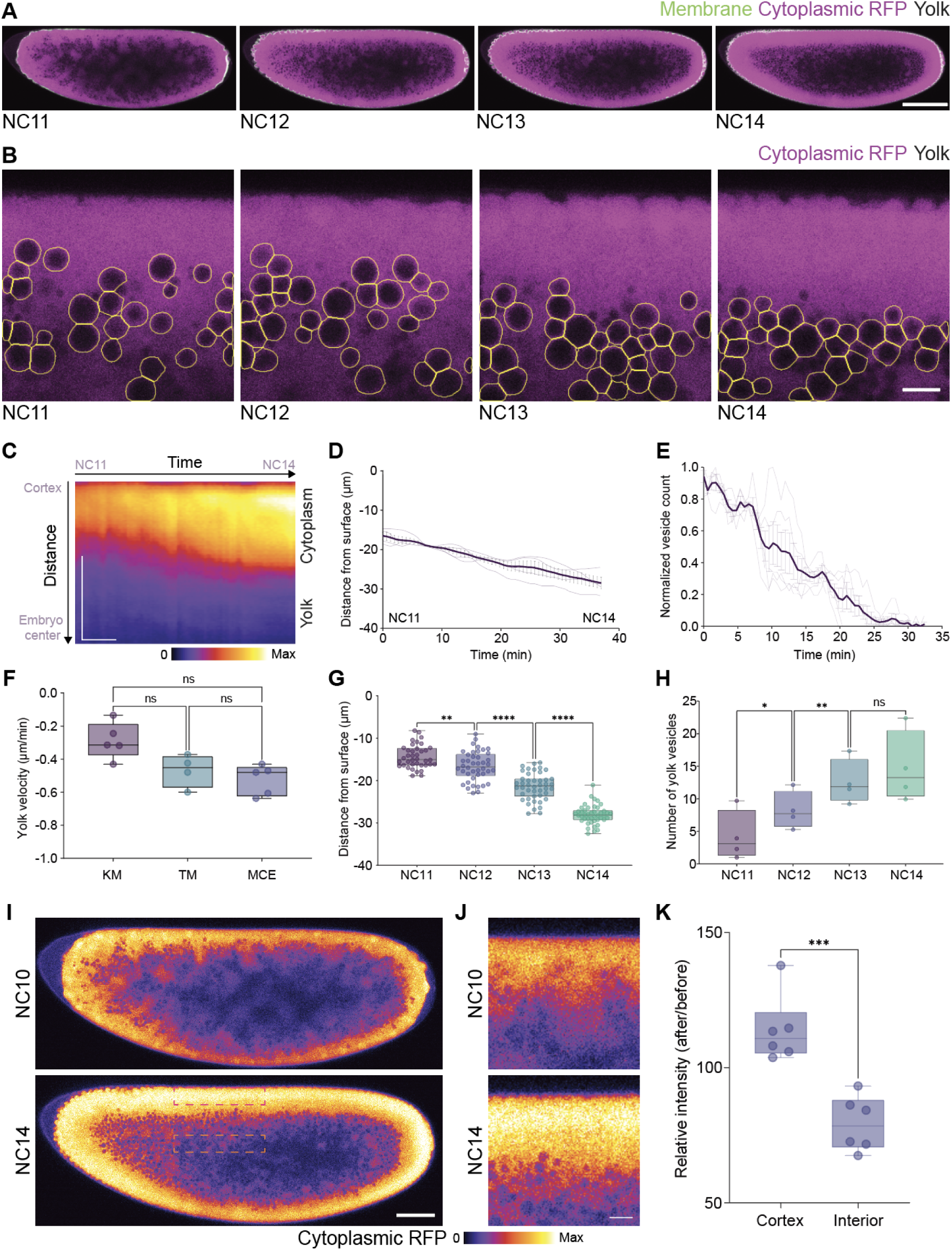
Progressive movement of the yolk vesicles from the embryo cortex to the center during NC11-14 in the *Drosophila* syncytial blastoderm. (A) Representative snapshots of live embryo imaging showing mRFP (cytoplasm, magenta), GFP-CAAX (cell membrane, green), and the yolk vesicles (unlabeled, dark regions), demonstrating gradual inward movement of the yolk vesicles from the embryo surface toward the center from NC11 to NC14. Scale bar = 100 μm. (B) Higher magnification views of yolk vesicle dynamics from time-lapse imaging as in (A), with automated segmentation of the yolk vesicles outlined in yellow. Scale bar = 10 μm. (C) Kymograph derived from a cortex-to-center line profile, illustrating the temporal progression of yolk segregation from NC11 to NC14. Scale bar = 50 μm (vertical), 10 min (horizontal). (D) Quantitative analysis of the yolk-cytoplasm boundary from NC11 to NC14. The graph shows trajectories from multiple embryos (n = 4 embryos), using segmentation and processing of kymographs as the one shown in (C). Error bars are mean±SEM. (E) Normalized yolk vesicle count as a function of time from 3D time-lapse imaging (Figure S1H, S1I; n=5 embryos), demonstrating a progressive inward movement of the yolk vesicles toward the embryo center. Error bars are mean±SEM. (F) Quantification of yolk vesicle velocities based on three approaches: linear fit of the kymograph (“KM”, n=5 embryos), direct tracking of the yolk vesicles (“TM”, n=4 embryos), and fitting to a mass conservation equation (“MCE”, n=5 embryos). One-way ANOVA with Bonferroni’s multiple comparisons test. Data presented here and henceforth as box plots show the median (center line), interquartile range (box), and minimum/maximum values (whiskers). (G) Measurements of the outermost yolk vesicles to the embryo surface (referred to as “yolk-to-surface distance” henceforth) from NC11 to NC14 (n=168 vesicles from NC11-14). Data show a progressive inward movement with a mean distance of 14.4 ± 0.44 μm (NC11), 16.6 ± 0.54 μm (NC12), 21.4 ± 0.48 μm (NC13), and 28.0 ± 0.34 μm (NC14). One-way ANOVA with Bonferroni’s multiple comparisons test. (H) Quantification of yolk density in the embryo center from NC11 to NC14, showing progressive densification of the yolk vesicles (n=4 embryos): 4.2 ± 1.9 (NC11), 8.2 ± 1.4 (NC12), 12.5 ± 1.7 (NC13), and 14.7 ± 2.7 (NC14) vesicles per 1000 μm^2^. One-way ANOVA with Bonferroni’s multiple comparisons test. (I, J) Cytoplasmic RFP distribution before (NC10) and after (NC14) yolk segregation, shown as a full embryo view (J) and in a higher magnification view to highlight the embryo cortex. The dotted boxes in the bottom panel in (I) show regions in the embryo cortex and interior, where the intensity was measured for (K). Scale bar = 50 μm (I), 10 μm(J). (K) Box plot showing ratios of RFP intensity after and before yolk segregation at the embryo cortex and interior. Welch’s t test (n = 6 embryos). Annotations for the statistical significance tests are as follows and used across all figures: ns, not significant, *p < 0.1, **p < 0.01, ***p < 0.001, ****p < 0.0001.

Accompanying the inward movement and densification of the yolk vesicles in the embryo center, we found an increase of cytoplasmic RFP concentration at the embryo cortex (Figure 1I-K,Movie S3). In sum, our data reveal a reciprocal movement of yolk vesicles and cytoplasm from NC11 to NC14 in the final stages of the *Drosophila* syncytial blastoderm. We term this process "yolk segregation and compaction".

### The progressive movement of the yolk vesicles correlates with microtubule axial expansion

Previous studies have implicated cell cycle dynamics and its associated cytoskeletal apparatus in the process of yolk-cytoplasm segregation in several organisms including *Drosophila* ^7,18,20^. Nuclei arrive at the embryo cortex by NC10 in *Drosophila*, bringing two major cytoskeletal systems to the cortex ^21^: 1) actin, which forms a cap-like cortical structure at the embryo surface above each nucleus; 2) microtubules, nucleated from the centrosomes associated with the nucleus. From NC10 onward, we detected a high degree of correlation between the planar movement of nuclei ^22^ and that of the yolk vesicles in the plane just underneath the nuclei (Figure S1J; analyzed via *en face* 3D imaging and segmentation, as shown in Figure S1G, H), with yolk vesicle velocity tracking closely that of the nucleus (Figure S1K). The correlative movement of the yolk vesicles and cortical nuclei raised the possibility that nuclei-associated cytoskeletal systems drive yolk-cytoplasm segregation. Thus, we asked which of these two events of cytoskeletal remodeling correlate with yolk vesicle movement.

Imaging of an F-actin marker (LifeAct-mCherry or Utrophin-mCherry) reveals cortically enriched and spatially confined F-actin primarily within 10 μm from the embryo surface (Figure 2A, B). Quantitative analysis shows that the spatial profile and the cortical intensity of F-actin remains largely unchanged from NC11 to NC14 (Figure 2D, E, G, H). This contrasts with the yolk vesicles that become progressively distanced from the embryo surface, producing a gap that increases from ∼5 μm to nearly 20 μm. (Figure 2A, B, D, E, Movie S4: LifeAct-mCherry, Utrophin-mCherry).

**Figure 2:**
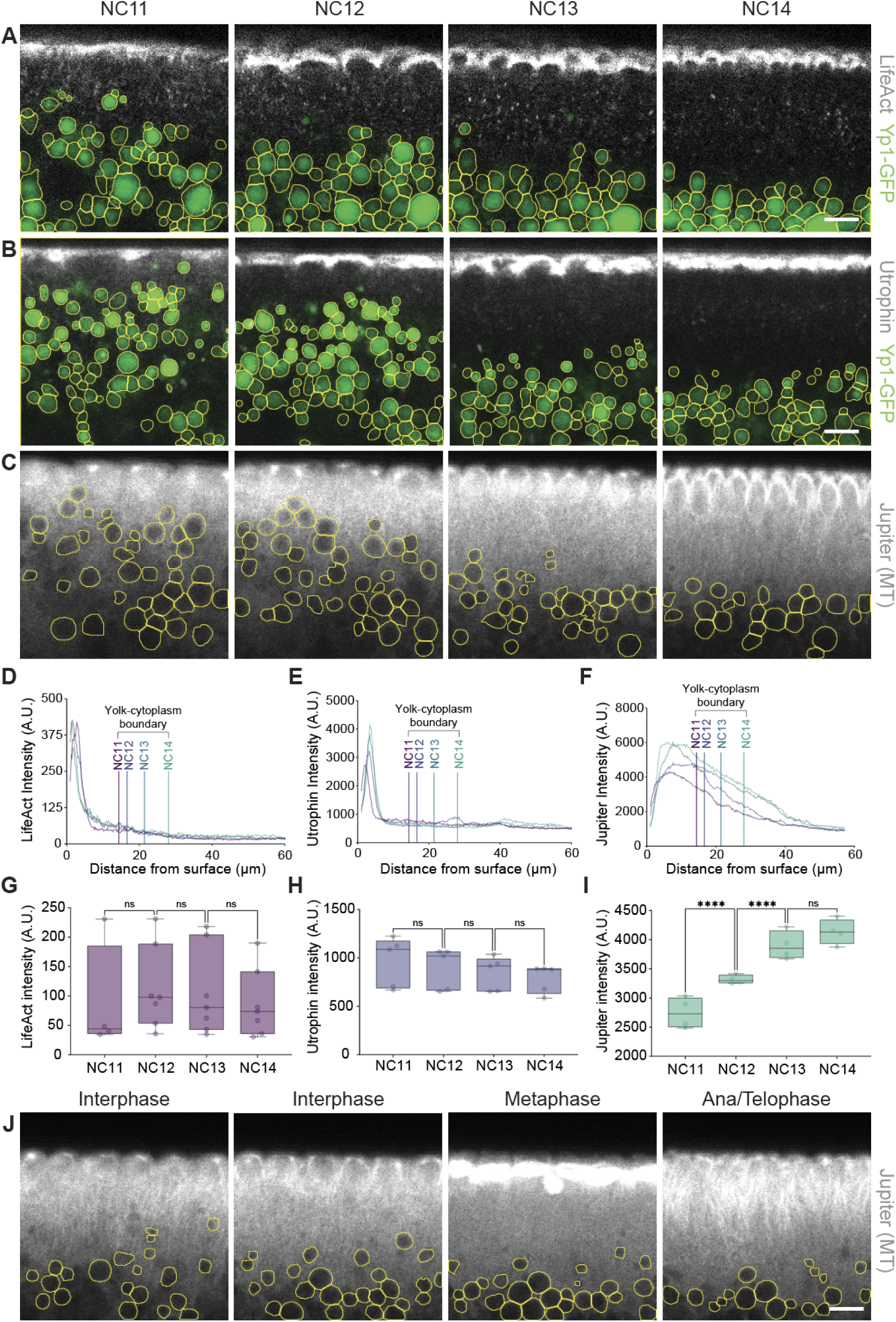
The yolk-cytoplasmic boundaries correlate with microtubule dynamics at the embryo cortex, but not that of actin. (A-C) Time-lapse imaging of LifeAct-mCherry (A, gray) or Utrophin-mCherry (B, gray) for F-actin, or Jupiter-GFP for MTs, in conjunction with yolk vesicle visualization using Yp1-GFP imaging (A, B, green), or negative labeling (C, exclusion of Jupiter-GFP) from NC11 to NC14. Segmentation of the yolk vesicles is shown with yellow outlines. (D-I) Quantification of F-actin (D, G, LifeAct, n=8 embryos; E, H, Utrophin, n=5 embryos) or microtubule (F, I, Jupiter, n=4 embryos) intensities. The intensities are represented as a function of distance (D-F) shown as line intensity profiles from the embryo surface to the center with annotated mean yolk-cytoplasm boundaries based on the measurements shown in Figure 1G. Intensities shown as a function of time from NC11 to NC14 (G-I) shown as box plots of intensity measurements taken from the region between the nuclei and the yolk. One-way ANOVA, mixed effects model, with Holm-Šídák’s multiple comparisons test. (J) Stage-dependent microtubule dynamics, as visualized by Jupiter-GFP, in relation to yolk vesicle distribution shown as segmented yolk vesicles (yellow outlines) within NC12. Scale bar = 10 μm.

These data did not reveal an association between actin and yolk vesicle dynamics. We do note, however, that in fixed embryos phalloidin labeling detects a low intensity, graded distribution of F-actin not detectable with live F-actin probes that extend from the yolk-free cortex to the yolk-rich interior (Figure S2A; see Discussion). In contrast, the centrosomal MTs, visualized by GFP-tagged microtubule binding protein Jupiter or by staining of alpha-tubulin, extended a much broader region from the embryo surface into the embryo center (Figure 2C, S2B).

Importantly, the vertical reach and the cortical intensity of MTs increases as NC progresses, tracking closely the inward movement of the yolk-cytoplasm boundaries (Figure 2F, I, Movie S5). We also observed progressive inward movement of the yolk vesicles within a NC as the microtubule architecture cycles through mitotic phases (Figure 2J). Overall, these data reveal a correlation between inward movement of the yolk and the spatial expansion of centrosomal microtubules across NCs.

### Destabilization of microtubule plus ends perturbs yolk-cytoplasm segregation

To functionally test whether yolk vesicle mobility requires microtubules, we used RNAi to knockdown EB1, a microtubule end-binding protein that localizes to, stabilizes, and promotes microtubule plus-ends growth ^23^. EB1 RNAi embryos generated aberrant microtubule structures (Figure 3A), consistent with previous reports ^24^. From NC11 to NC14, negative labeling of the yolk reveals aberrant, inhomogeneous yolk segregation, producing a jagged contour of yolk-cytoplasm boundary (Figure 3B). Quantitative analyses show a decreased displacement of the yolk towards the embryo center, perturbed cortical clearance of the yolk vesicles, and reduced yolk vesicle velocity, as compared to the control embryo (Figure 3C-F, S3A-B, Movie S6). Together, these observations suggest that microtubule stability and plus-end growth are required for yolk-cytoplasm segregation.

**Figure 3:**
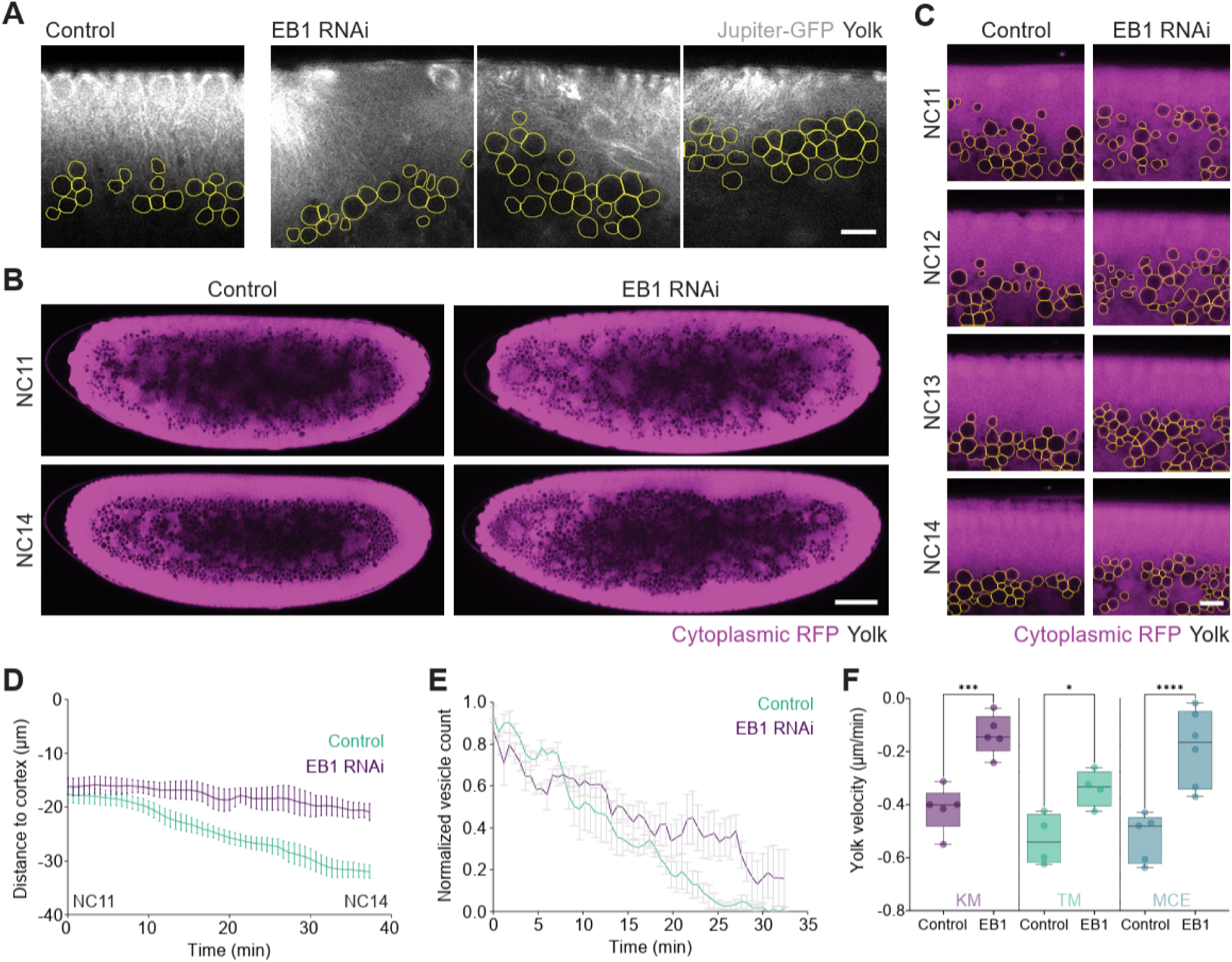
Microtubule growth and stability is required for proper yolk segregation. (A) Images of a control and three representative EB1 RNAi embryos showing normal (control) and perturbed (EB1 RNAi) centrosomal microtubule structures at the NC13 embryo cortex, as visualized by Jupiter-GFP. The yolk vesicles can be seen as negatively labeled circular objects that are segmented and outlined with yellow contours. Scale bar = 10 μm. (B, C) Representative time-lapse images of control and EB1 RNAi embryos with cytoplasmic RFP (magenta) showing disturbed yolk segregation in EB1 RNAi embryo, where the yolk-cytoplasm boundary appears jagged at the embryo scale (B), and the in-ward mobility of the yolk vesicles is reduced (C; the yolk vesicles are outlined in yellow). Scale bars = 50 μm (B) and 10 μm (C). (D) Kymograph derived quantitative analysis of the yolk-to-cortex distance from NC11 to NC14 (n=5 embryos for both). Error bars are mean±SEM. (E) Normalized yolk vesicle count as a function of time at the outermost 33 μm of embryo cortex (Control: n=5 embryos; EB1 RNAi: n=6 embryos). Error bars are mean±SEM. (F) Yolk velocities as estimated by the three methods (“KM”, “TM” and “MCE”) outlined above. Number of embryos [control, EB1 RNAi] = [5, 5] (“KM”), [4, 4] (“TM”), [5, 6] (“MCE”). Control “MCE” data are the same as those used in Figure 1F. One-way ANOVA with Bonferroni’s multiple comparisons test.

### Gnu depletion decouples nuclear dynamics from yolk-cytoplasm segregation

Actin and microtubule networks are known to be interdependent, while perturbing either one of them can disrupt cortical nuclear migration and induce nuclear fall out from the cortex ^25,26,27^. To causally assess the role of cortical cytoskeletal networks in yolk segregation, it would thus be essential to use a genetic background in which the cortical cytoskeletal systems can be established despite the absence of cortical nuclear migration. To do so, we exploited the possibility of using embryos lacking *gnu* to decouple cortical cytoskeletal networks from the nuclear dynamics. Gnu is a component of the PanGu complex, which ensures adequate cyclin B levels during early mitosis ^28^. In *gnu* deficient embryos, DNA replication proceeds normally, but the replicated chromosomes fail to segregate, causing the replicated DNA to aggregate into a small number of giant nuclei, leading to the loss of an evenly distributed layer of cortical nuclei ^29,30^. Importantly, centrosomes continue to duplicate independently in these embryos and eventually migrate to the cortex, where they produce extensive microtubule networks, despite the disruptions of nuclear divisions ^29,30^. In the *gnu* RNAi embryos, we observed unsegregated DNA masses, confirming previous reports^29,30^ (Figure 4A, Movie S7; visualized by H2Av-RFP). In contrast, the centrosomes – visualized by Jupiter-GFP – continued to divide, and moved to the cortex, despite having been uncoupled from the nucleus (Figure 4A, B; Movie S7).

**Figure 4:**
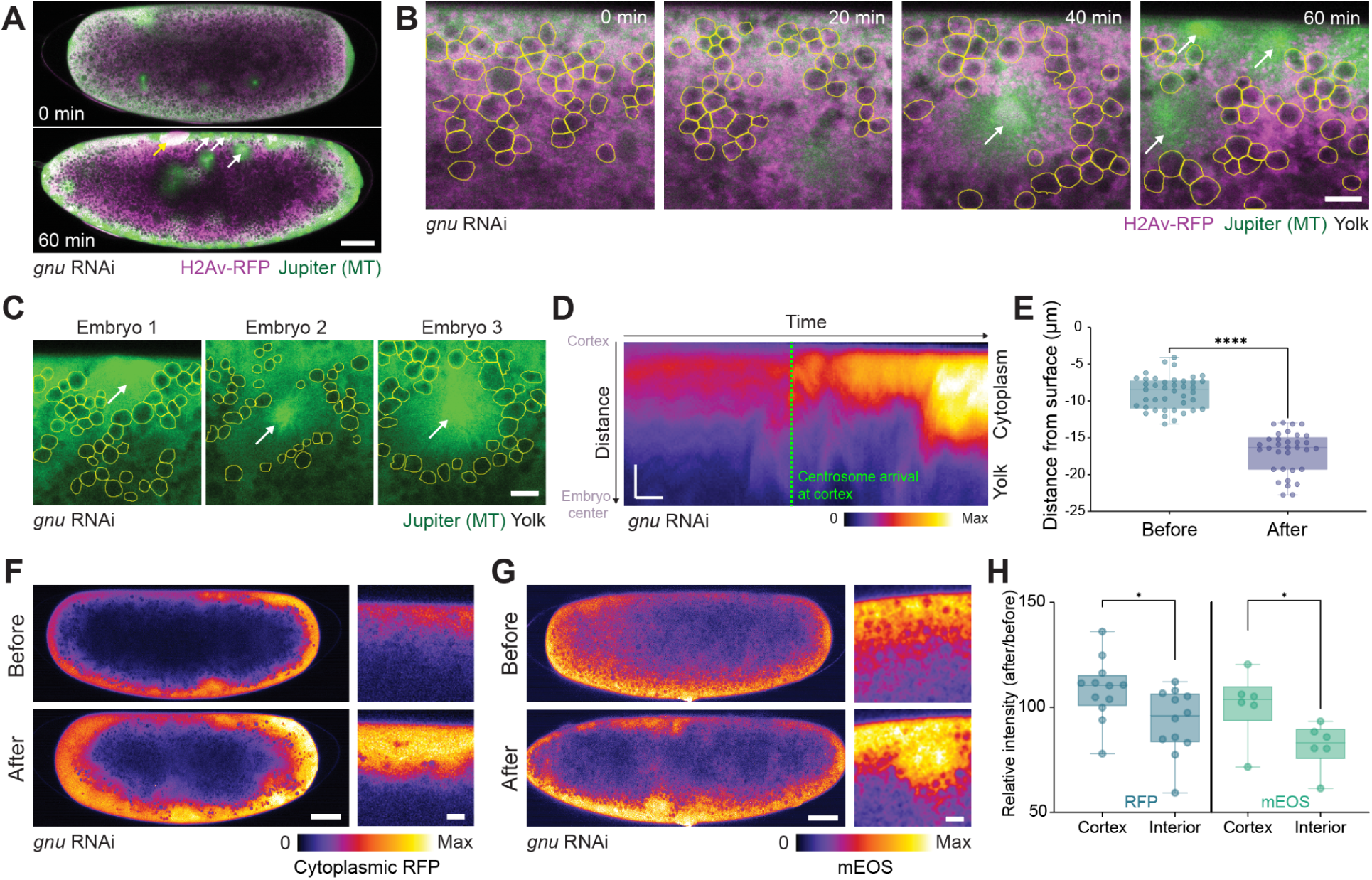
Centrosomally organized microtubules in Gnu depleted embryos drive yolk segregation. (A) Representative snapshots from live imaging of embryos expressing H2Av-RFP (nucleus, magenta) and Jupiter-GFP (MTs, green) showing centrosome (white arrows) segregation and migration to the embryo surface, despite defective chromosome segregation and the formation of giant nuclei (yellow arrow). Note that H2Av is known to be sequestered at the lipid droplets, in addition to localizing to the nucleus, thus labeling the embryo interior ^31^. Scale bar = 50 μm. (B) High magnification view of (A) showing the arrival of centrosomes (white arrows) at the cortex, in relation to the yolk vesicles (segmented yellow outlines), revealing the clearance of the yolk from centrosomal asters, forming yolk-free zones. Since the *gnu* RNAi embryo cannot be staged based on the NCs, we defined t=0 min as time before cortical migration of the centrosomes. Scale bar = 10 μm. (C) Three representative *gnu* RNAi embryos showing clearance of yolk vesicles in the regions surrounding centrosomes (Jupiter, green) either at the cortex (Embryo 1) or in the embryo interior (Embryos 2, 3). (D) A kymograph derived from the cortex-to-center line profile showing the temporal correlation between the arrival of centrosomes and the onset of yolk segregation and cortical clearance. The green dotted line represents the timing of centrosome arrival. Scale bar = 10 μm (horizontal), 10 min (vertical). (E) Quantification of yolk vesicle distances from the embryo surface before and after centrosome arrival (n = 41,33 vesicles, from at least 3 embryos). Data show significant increase in yolk-surface distance following centrosome positioning. Mann-Whitney test. (F-G) Cytoplasmic mRFP (F) and photoactivated mEOS (G) distribution before and after yolk segregation, shown in a full embryo view and in higher magnification to highlight the embryo cortex. Scale bar = 50 μm (full embryo), 10 μm (higher magnification). (H) Box plot showing significant enrichment of RFP and mEOS in the embryo cortex, highlighting enrichment after yolk segregation in the cortex, and the lack of such enrichment in the embryo interior. Welch’s t test (n = 12 for mRFP; 6 for mEOS).

Strikingly, the centrosomes created yolk-free zones in their immediate vicinity either at the embryo cortex (Figure 4C: Embryo1) or in the embryo interior (Figure 4C: Embryos 2, 3), suggesting that centrosomes can cause yolk clearance, even in the absence of the nuclei. Kymograph analysis further confirmed correlation between centrosome arrival and increased region of yolk clearance at the cortex (Figure 4D, E). As further evidence for the *gnu* RNAi embryos to be suitable for investigating yolk segregation, following the arrival of centrosomes to the cortex, the cortical cytoplasmic RFP concentration indeed increased, mirroring that of control embryos (Figure 4F, 4H, Movie S3, as compared to Figure 1I-K). To ensure such an increase is due to the concentration of existing cytoplasmic proteins, but not the newly synthesized or fluorescently matured proteins, we performed a pulse-chase experiment with the photoconvertible mEOS fluorescent protein and found that the photoconverted pool of mEOS protein also became concentrated at the cortex after cortical migration and yolk segregation (Figure 4G, 4H, Movie S3). These data demonstrated that there was indeed concentration of the cytoplasm in the *gnu* RNAi embryos during yolk segregation. Taken together, the hallmarks of yolk segregation can be observed in *gnu* RNAi and are decoupled from cortical nuclear migration, validating its use as an experimental system to investigate the role of cytoskeleton in yolk-cytoplasm segregation.

### Yolk-cytoplasm segregation in the *gnu* RNAi embryos depends upon microtubules, but not actin

We then proceeded to use a pharmacological approach to functionally investigate the contribution of microtubule and actin networks to yolk-cytoplasm segregation in the *gnu* RNAi embryos. We first injected colcemid to depolymerize microtubules. In embryos in which the centrosomes had not yet divided and migrated to the cortex, colcemid severely impaired the clearance of the yolk vesicles from the cortex (Figure 5A-C Movie S8; “Colcemid early”). The yolk-to-surface distance remained unchanged 40 minutes after the injection, contrasting with water (vehicle for colcemid) injected control embryos (Figure 5A-C,”Water”). Similarly, in embryos in which centrosomal division and cortical migration had occurred, colcemid treatment prevented any further increase of yolk-to-surface distance (Figure 5A-C, Movie S8; “Colcemid late”). In sum, the vertical mobility of the yolk vesicles was severely impaired in the colcemid-injected embryos, irrespective of the timing of injection. It is worthwhile noting that the anterior-posterior oscillations of the yolk, a process known to be driven by actomyosin-dependent cytoplasmic streaming ^32^, continued to occur, despite the depolymerization of microtubules (Movie S8).

**Figure 5:**
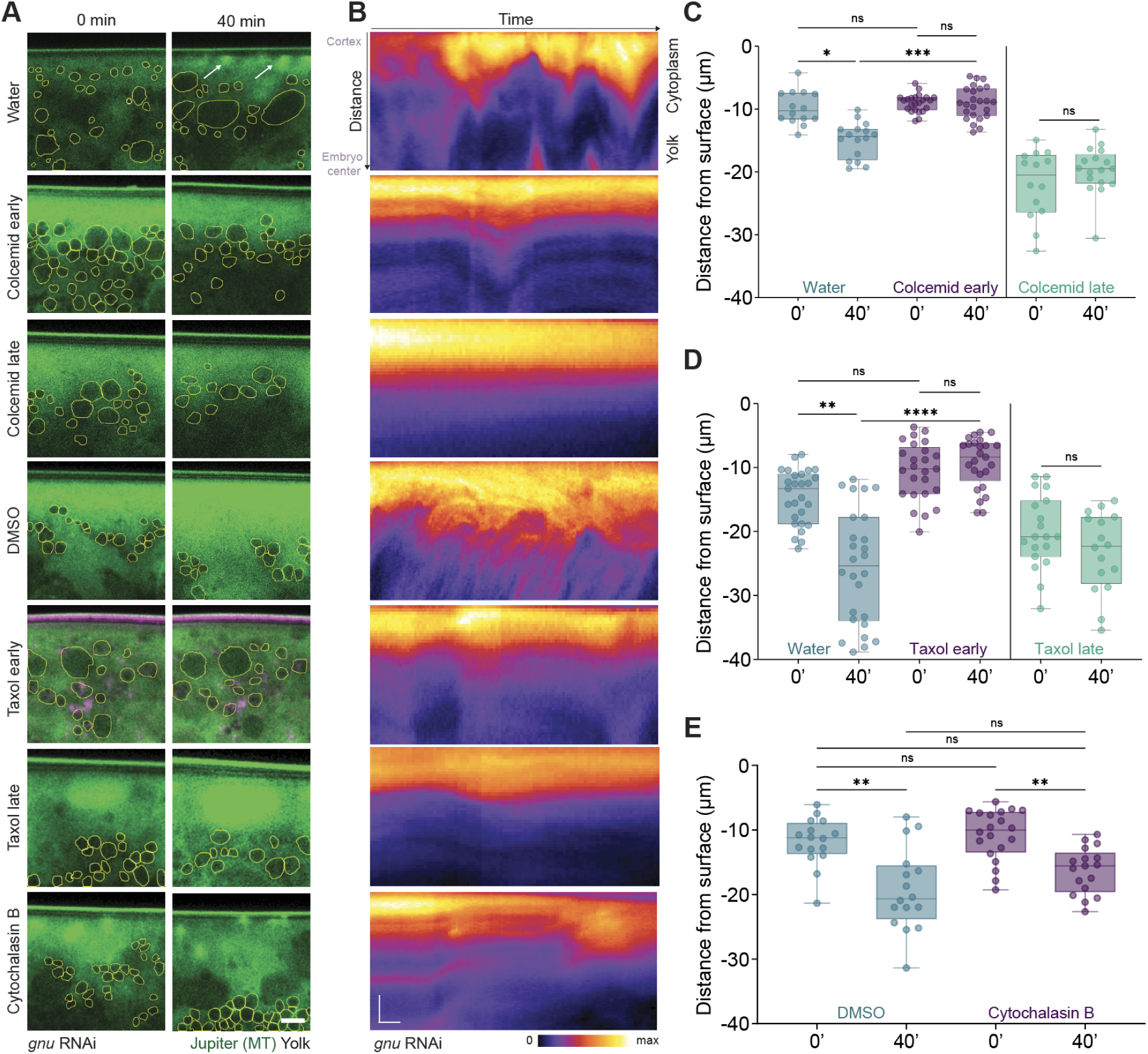
Inward mobility of the yolk vesicles require dynamic microtubules, but not actin. (A) Pharmacological inhibitor injections into the *gnu* RNAi embryos labelled with Jupiter-GFP, shown as snapshots at the time of injection (0 min) and 40 min later. (B) Kymographs corresponding to images shown in (A) for visualization of the distribution of yolk and cytoplasm as a function of time. As compared to water (vehicle control) injected embryos, where the yolk-free zone at the embryo cortex widened, in colcemid injected embryos (Colcemid early, injection performed before centrosomal arrival; Colcemid late, injection performed after centrosome arrival), the yolk-free zone at the cortex showed minimal changes. Similar results were obtained based on a comparison between DMSO (vehicle control) injected embryos, where the yolk-free zone widened, and Taxol injected embryos, either injected before centrosomal arrival (Taxol early) or after centrosome arrival (Taxol late), where the yolk-free zone at the cortex showed minimal changes. In contrast, Cytochalasin B injected embryos showed widening of the yolk-free zone at the cortex. Scale bars = 10 μm for the images; 10 μm (horizontal), 10 min (vertical) for the kymographs. (C) Quantification of yolk vesicle distances from surface in early colcemid, late colcemid, and water-injected control embryos (n =23, 24,14,16,14,17 vesicles, from at least 3 embryos and in the order of representation). Kruskal-Wallis (non-parametric) test. (D) Similar experimental design and analysis as (C) with taxol treatment to stabilize microtubules. For the quantification, in (D), n =25, 25, 27, 26,19,16 vesicles, from at least 3 embryos, in the order of representation. Kruskal-Wallis (non-parametric) test. (E) The yolk vesicles move inward in Cytochalasin B treated embryos. Injections were performed only in the early stage before centrosome arrival, (n =16,16, 20,17 vesicles, from at least 3 embryos, in the order of representation). Kruskal-Wallis (non-parametric) test.

We next injected taxol into the *gnu* RNAi embryos to inhibit MT turnover. When injected early, before centrosome arrival at the cortex, taxol blocked yolk segregation, similar to colcemid (Figure 5A, B, D, Movie S8; “Taxol early”). In later-stage embryos where yolk segregation had already occurred at the cortex following the arrival of centrosomes, taxol prevented yolk from moving further towards the embryo center, again mirroring the effect of colcemid injection (Figure 5A, B, D, Movie S8; “Taxol late”). These results contrast with those of the carrier (i.e. DMSO) injected embryos, where yolk-to-surface distance increases substantially, 40 minutes after the injection (Figure 5A, B, D). Together, these observations suggest that microtubule dynamics is necessary for yolk-cytoplasm segregation and corroborate our data on EB1 RNAi.

In contrast, depolymerizing actin using cytochalasin B did not prevent the yolk vesicles from moving, while the centrosomes can be seen dividing and clearing the yolk vesicles away from their immediate vicinity (Figure 5A, B, E, Movie S8; “Cytochalasin B”). Importantly, cytochalasin B injection indeed eliminated the anterior-posterior oscillations of the yolk vesicles, which, as mentioned above, require actomyosin contractility (Movie S8) ^32^. Loss of these oscillatory movements caused the centrosomes to cluster in localized regions, rather than spreading along the anterior-posterior axis of the embryo. Interestingly, we could see that clustered centrosomes create enlarged zones of yolk vesicle clearance, providing additional support that microtubules clear away the yolk vesicles.

In sum, pharmacological perturbations of microtubules and actin in the *gnu* RNAi embryos show that the polymerization/depolymerization dynamics of the centrosome-derived microtubule networks is necessary to drive yolk-cytoplasm segregation. In contrast, depolymerization of actin networks does not prevent yolk clearance from the cortex and from the vicinity of the centrosomes. We conclude that active MT dynamics drives yolk-cytoplasm segregation.

### EB1 comets come in apparent contact with and displace the yolk vesicles

We further characterized microtubule-yolk vesicle interactions in the *gnu* RNAi embryos to gain insights into how microtubule dynamics drives yolk-cytoplasm segregation. We imaged the yolk vesicles alongside growing microtubule plus-ends using EB1-GFP (Figure 6A, Movie S9,10). Imaging *en face* allowed us to identify regions of the embryo cortex where the centrosomes had newly arrived. EB1 comets can be seen emanating from these centrosomes, producing characteristic firework-like patterns of an outward growing aster (Figure 6A, green arrows). Notably, EB1 comets can be seen to come in apparent contact with the yolk vesicles (Figure 6A, yellow arrows). Some of these vesicles exhibited motions of moving beneath the imaging plane and ultimately vanished from the view (Figure 6A, yellow arrows, Embryo3). These observations suggest that the growing microtubule plus ends physically displace the yolk vesicles toward the center of the embryo.

**Figure 6:**
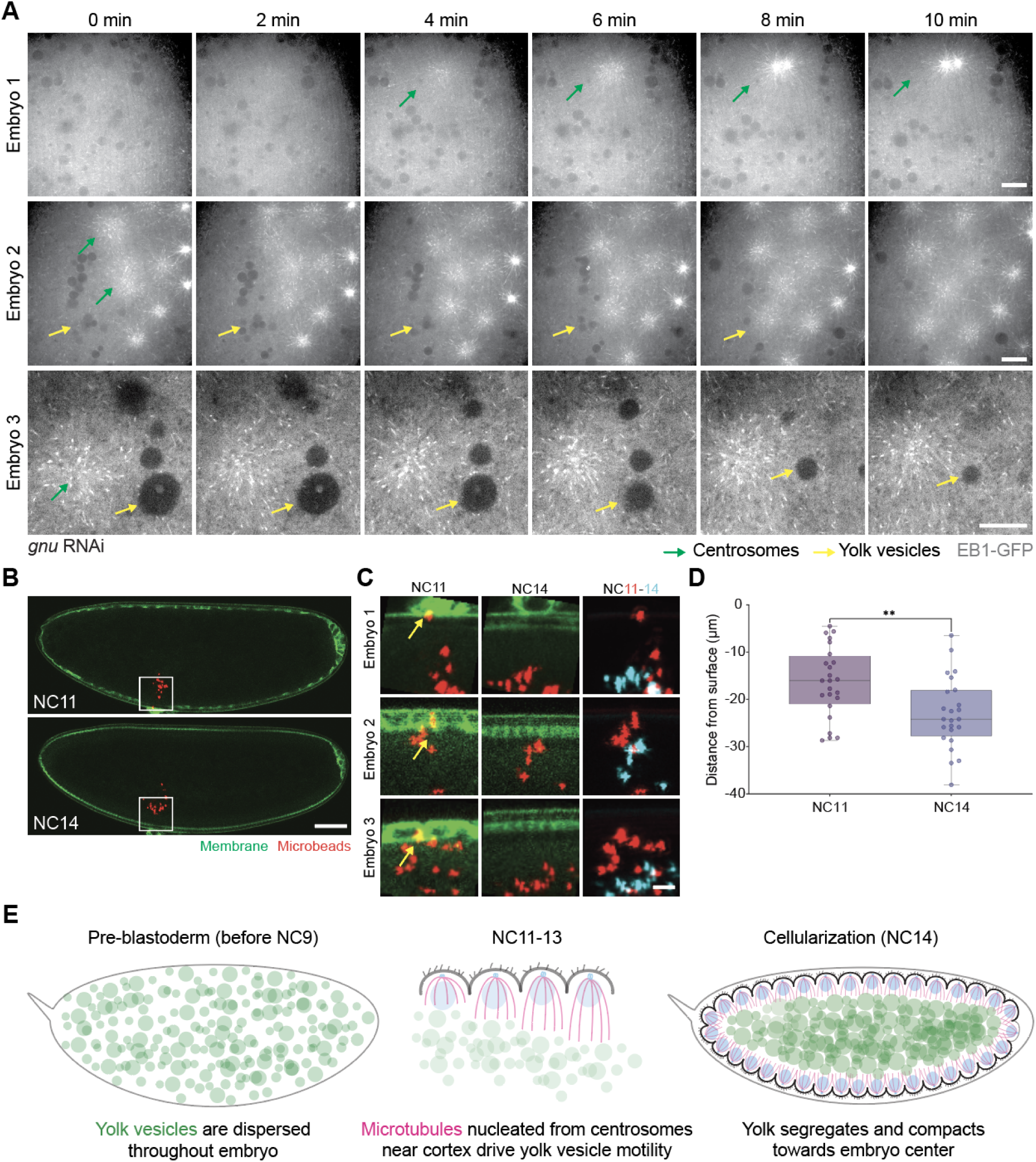
Evidence supporting microtubule plus end growth displaces the yolk via direct contact. (A) Time-lapse imaging of *the gnu* RNAi embryos expressing EB1-GFP (microtubule plus-end marker). Three representative embryos demonstrate yolk segregation patterns upon centrosomal arrival. Embryo 1 shows an increase of the yolk clearance region following centrosome arrival (green arrows). Embryo 2 shows the expanding yolk free zone around the growing centrosomal asters (green arrows) that is accompanied by the disappearance of the yolk vesicles (yellow arrows) as they move towards the embryo center. Embryo 3 provides a closeup view of the close apposition of an extensile microtubule aster (green arrows) with a few yolk vesicles (yellow arrows) that are gradually displaced and become submerged underneath the imaging plane. Time indicates minutes elapsed from the start of imaging. Scale bar = 10 μm. (B) Representative images of control embryos at NC11 (top) and NC14 (bottom), following the injection of PEG-passivated fluorescent beads (1 µm diameter, red). The membrane is marked with GFP-CAAX (green). White boxes indicate regions magnified in (C). Scale bar = 50 μm. (C) Higher magnification view showing the injected beads distribution in three representative embryos at NC11 and NC14. Left column: NC11 embryos with beads positioned near the embryo surface (yellow arrows). Middle column: NC14 embryos showing beads displaced toward the embryo interior. Right column: superimposition of NC11 (cyan) and NC14 (red) images, showing beads displacement. Scale bar = 10 μm. (D) A box plot for the quantification of beads distance from the embryo surface at NC11 and NC14, with beads positioned at significantly deeper locations at NC14, as compared to NC11. Each data point represents an individual bead (n = 24 beads per condition from 5 embryos). Mann-Whitney test. (E) Schematic representation of the model for yolk segregation in the *Drosophila* embryo: Dispersed yolk vesicles are segregated from the embryo cortex, driven by the dynamic growth of cortical MT networks.

### Movement of the injected inert beads mirrors that of the yolk vesicles

We reason that if the growing microtubule plus ends push and displace the yolk vesicles, non-biological, inert objects of a similar size, when introduced into the embryo, could be subject to the same physical force and be similarly displaced towards the embryo center. To test this possibility, we injected fluorescently labeled microbeads with a diameter of 1 μm into the embryos. We passivated these microbeads with polyethylene glycol (PEG) prior to injection to reduce their non-specific binding to cellular components. When positioned near the embryo surface, the injected beads gradually move inward towards the embryo center from NC11 to NC14 (Figure 6B,C, Movie S11). Beads initially located in the yolk region showed variable behaviors, including rare cases where they moved toward the cortex (Figure 6B,C). Quantitative analysis, however, showed that overall the beads moved from an averaged position of ∼15 μm from the embryo surface at NC11 to ∼25 μm at NC 14, thus in quantitative agreement with yolk vesicle displacement (Figure 6D). These data thus provide support for the existence of non-selective, directional, and potentially direct force that can displace micron-sized spherical objects towards the embryo center.

## Discussion

Yolk-cytoplasm segregation is one of the earliest organizational and spatial partitioning events in the embryo of the oviparous animals, yet the underlying mechanisms have remained poorly understood. Our study reveals that in the syncytial *Drosophila* embryo, yolk segregation is primarily driven by microtubule dynamics (Figure 6E). Using quantitative live imaging, genetic perturbations, and pharmacological inhibitors, we show that the yolk vesicles move progressively inward during NC 11–14 in tight coordination with the expanding microtubule networks that increases its spatial reach over nuclear cycles. The filamentous actin structures, by contrast, remain predominantly cortical, and do not evolve in their spatial distribution during this period. The most compelling evidence comes from the *gnu* RNAi embryos, where the microtubule asters that are nucleated by the centrosome are decoupled from the syncytial nuclei, allowing us to use a pharmacological approach to establish causality. Specifically, the microtubule asters create local yolk-free zones, cortically clearing the yolk vesicles, while injection of either colcemid or taxol that disrupts the MT dynamics, block yolk clearance and yolk vesicle mobility. In contrast, actin depolymerization does not alter these behaviors. These data suggest a model whereby dynamic MT polymerization drives the displacement of the yolk vesicles, a process we successfully recapitulate with injected, passivated fluorescent microbeads.

### Mechanistic basis of MT based yolk vesicle segregation

MTs form rigid polymers that can move and deform cellular structures including nuclei ^33,34,35,36^, in addition to their better known function as cellular highways on which the organelles as cargoes are translocated by the motors ^37^. MT asters are known to centre centrosomes in large cells, whereby MT polymerization exerts pushing forces against the cellular cortex to position the centrosomes ^38^. Our data are better aligned with the interpretation that growing microtubule polymers exert direct pushing forces on the yolk vesicles, rather than translocating them via motor-driven mobility. Several independent lines of evidence support this interpretation. First, despite that taxol is known to preserve motor-driven mobility ^39,40^, injection of taxol into the *gnu* RNAi embryo prevents yolk segregation. Second, measured yolk displacement rate (∼0.3 μm/min) is in agreement with MT growth rate (1-7 μm/min), while nearly two orders of magnitude slower than kinesin-driven cargo velocity (0.1–0.3 μm/s) ^41,42, 43–45^. Finally, passivated, inert microbeads of a comparable size are displaced towards the embryo center in a manner similar to the yolk vesicles, corroborating the plausibility of direct, physical displacement, rather than motor-driven mobility. Importantly, our *in vivo* observations of microbead displacement are in line with the previous *in vitro* demonstrations that microtubule asters can physically repel and clear the yolk vesicles ^30^. Together, these observations suggest a flashing ratchet-type model ^46^, in which polymerizing microtubule arrays displace the yolk vesicles, thereby progressively compacting yolk toward the embryo interior.

### Unresolved role of actin

Although depolymerization of the filamentous actin network using Cytochalasin B did not have any discernible effect on yolk segregation in the *gnu* RNAi embryos, we were nonetheless intrigued by the meshwork-like, graded pattern of bulk actin distribution in the embryo interior ranging from the subcortical to the yolk-rich regions of the embryo, as revealed by the phalloidin staining (Figure S2A). It remains to be seen whether these actin structures play a role in yolk segregation that our assay was not able to detect, or whether they are functionally associated with any other processes of yolk spatial organization. Future studies should employ more specific perturbations of actin function combined with higher-resolution imaging.

### Functional significance and cytoplasmic organization

Yolk segregation is not simply a repositioning of the yolk vesicles towards the embryo center, but also the corresponding repartitioning of the soluble cytosolic fraction towards the embryo cortex. As was observed with cytoplasmic mRFP in control and the *gnu* RNAi embryos, and further verified with the photoconvertible mEOS in the *gnu* RNAi embryos, the densification of yolk materials in the embryo center is accompanied by the reciprocal enrichment of cytoplasmic proteins near the embryo surface. While it remains to be seen how yolk compaction might be directly or indirectly involved, the concentration of cytoplasm could potentially have profound functional significance for early embryonic development. Since most cytoplasmic components are maternally deposited prior to the onset of zygotic genome activation, which occurs during NC14 ^21,47^, concentrating maternal deposits may represent an adaptive feature whereby concentrating cytoplasmic substances, such as ribosomes, metabolic enzymes, and developmental regulators, could promote biochemical efficiency while conserving maternal resources.

### Evolutionary perspectives on yolk segregation mechanisms

The apparent divergence between *Drosophila* and zebrafish yolk segregation mechanisms raises an intriguing question of whether each strategy may be an evolutionary adaptation to a specific mode of early embryonic development. In zebrafish, yolk segregation occurs in the one-cell stage embryo, preceding the first embryonic cleavage, and thus cannot rely on the operation of mitotic spindles ^7^. In contrast, *Drosophila* yolk segregation trails behind the initial nuclear mitotic cycles that take place deep within the embryo and does not commence until the nuclei migrate out to the embryo cortex. That yolk displacement is temporally coupled to the cortical translocation of the mitotic machinery, may have facilitated the co-option of the pre-existing microtubule networks into the process of yolk segregation. These considerations invoke the concept of heterochrony in evolutionary biology ^48^, where changes in the developmental timetable can be found to be associated with – and possibly causally linked to – distinct morphological outcomes or divergent morphogenetic mechanisms.

To further explore this conceptual idea, it would be of particular interest to examine other species, in which the early embryonic mitoses precede or occur concurrently with yolk segregation, e.g. the *Xenopus*, *C. elegans*, and *Ascidian* embryos. In fact, recent work in the *C. elegans* resonates with our findings of a microtubule-based mechanism for yolk segregation^20^. Specifically, it was shown that the meiotic spindle of the oocyte is indirectly pushed outward to the cortex by a volume exclusion mechanism which depends on the kinesin motor based inward translocation of the yolk granules and mitochondria. While the polarized, plus-end-in-minus-end-out microtubule network organization in the *C. elegans* oocyte mirrors that in the early *Drosophila* embryo after nuclear cortical migration, the involvement of a motor-cargo based mechanism for yolk-cytoplasms segregation is distinct, raising the question of whether the size of yolk granules (0.5-1 μm in *C. elegans* ^49^ versus 1-9 μm in *Drosophila*) may be the relevant parameter that sets them apart. In sum, the timing of yolk segregation relative to cell/nuclear division, alongside species-specific yolk composition and granule size, may shape the evolutionary trajectories selecting the cytoskeletal mechanism underlying yolk-cytoplasm segregation.

### Future directions

Several key questions emerge from this work that would warrant and could motivate further investigation. It remains to be seen whether there are any microtubule-associated proteins or motor proteins that are involved in yolk displacement in ways that evade our assessment via taxol perturbation, and whether as a feedback, microtubule polymerization rates differ between the yolk-rich and yolk-depleted zones. Comparative studies in dipterans with different egg sizes or nuclear densities would test whether microtubule-based segregation is broadly conserved or a *Drosophila*-specific solution. Finally, determining whether defects in yolk segregation may affect early embryonic morphogenesis, such as cellularization, gastrulation, or early patterning, will shed light on the developmental significance of this fundamental organizational process.

## Acknowledgements

We thank the Bloomington and Kyoto *Drosophila* Stock Centers for reagents; Makito Miyazaki, Yosuke Yamazaki from RIKEN-IMS for assistance with passivation of microbeads with PEG; Kobe BioImaging Facilities and Factory (KBiIF), Shigeo Hayashi for support; members of the Kondo, Hayashi, Obata and Yoo laboratories for discussions; Bipasha Dey, T.-Y. Huang for critical reading and comments on the manuscript; members of Rikhy, Wang and Lemke labs, especially Somya Madan, Anne Rosfelter, Nada Dogui, Steffen Lemke, Girish Kale, Verena Kaul, and members of the Transformative Grants-in-Aid (A) for critical discussions.

## Author contribution

S.T., A.N., M.M., R.R. conceived, designed and supervised the study. S.T. performed experiments. S.S and S.G. performed phalloidin staining and imaging. S.T, P.D and B.K analysed the data. R.R. and Y.C.W arranged for funding and critical feedback. S.T. and Y.C.W wrote the manuscript and all other authors contributed in editing the manuscript.

## List of movies

**Movie S1.** Time-lapse imaging of a syncytial *Drosophila* embryo expressing cytoplasmic mRFP, where the yolk vesicles appear as dark, RFP-excluding objects that progressively move away from the cortex toward the embryo interior during NC11–14.

**Movie S2.** Live imaging of a *Drosophila* embryo expressing Yp1-GFP, which directly labels the yolk vesicles and allows for visualization of their inward movement during syncytial nuclear cycles.

**Movie S3.** Live imaging of cytoplasmic mRFP in control and the *gnu* RNAi embryos, showing progressive enrichment of the cytoplasmic signal at the cortex and the concurrent yolk segregation toward the embryo interior, consistent with cytoplasm-yolk phase separation.

**Movie S4.** Live imaging of *Drosophila* embryos co-expressing LifeAct-mCherry and Yp1-GFP, or Utrophin-mCherry and Yp1-GFP, showing that the cortical and cytoplasmic actin network remains predominantly restricted to the embryo surface as the yolk vesicles progressively move inward.

**Movie S5.** Live imaging of a *Drosophila* embryo expressing Jupiter-GFP (microtubule marker) showing the expanding microtubule network, which displays a clear spatial anti-correlation with yolk vesicle distribution as NC progresses.

**Movie S6.** Time-lapse imaging of EB1 RNAi embryos expressing cytoplasmic mRFP, demonstrating aberrant yolk vesicle movement as compared to controls.

**Movie S7.** Live imaging of *gnu* RNAi embryos expressing H2Av-RFP and Jupiter-GFP, showing centrosome duplication and migration in the absence of nuclear division, accompanied by the formation of local yolk-free zones as the centrosomal asters expand near the cortex.

**Movie S8.** Time-lapse imaging of *gnu* RNAi embryos injected with water, colcemid, taxol, or cytochalasin B, demonstrating that disruption of microtubule dynamics — but not actin depolymerization — blocks yolk clearance. Note that the occurrence of the anterior-posterior oscillatory movement seen in colcemid and taxol injected embryos. This was due presumably to cytoplasmic flows driven by acto-myosin contraction and a previous report ^32^ suggested that such an oscillatory movement helps spread the cortical nuclei across the embryo cortex. Supporting these assessments, the oscillatory movement was abolished in cytochalasin B injected embryos, which may have contributed to the clustering of the centrosomes toward the anterior pole.

**Movie S9.** Live imaging of a *gnu* RNAi embryo expressing EB1-GFP, showing that the growing microtubule plus-ends emanate from the centrosomes that are arriving at the embryo cortex with EB1 comets coming in apparent contact with and displace the yolk vesicles.

**Movie S10.** A second representative *gnu* RNAi embryo expressing EB1-GFP, revealing the formation of aster-like, firework-patterned microtubule arrays emanating from the centrosomes, with their immediate vicinity showing clearance of the yolk vesicles

**Movie S11.** Time-lapse imaging of control embryos injected with PEG-passivated fluorescent microbeads (1 µm diameter), showing that these inert particles, when placed near the embryo surface, undergo inward displacement in a manner that closely resembles yolk vesicle movement.

## Materials and Methods

### Drosophila melanogaster Stocks and Maintenance

Flies were maintained at 25°C on standard cornmeal agar medium under 12-hour light/dark cycles. The following *Drosophila* stocks were used in this study:

| Reagent type<br>(species) or resource | Source or reference | Identifiers |
| --- | --- | --- |
| *D. melanogaster*: Oregon-R<br>(wild-type) | Bloomington Drosophila<br>Stock Center | BDSC: 6630; RRID:<br>BDSC_6630 |
| *D. melanogaster*:<br>mat-GAL4-VP16 | Bloomington Drosophila<br>Stock Center | BDSC: 7062; RRID:<br>BDSC_7062 |
| *D. melanogaster*: Jupiter::GFP | Bloomington Drosophila<br>Stock Center | BDSC: 5659; RRID:<br>BDSC_5659 |
| *D. melanogaster*: H2Av::RFP | Bloomington Drosophila<br>Stock Center | BDSC: 23650; RRID:<br>BDSC_23650 |
| *D. melanogaster*: ncd-EB1::GFP | Bloomington Drosophila<br>Stock Center | BDSC: 35512; RRID:<br>BDSC_35512 |
| *D. melanogaster*: UAS-EB1-RNAi | Bloomington Drosophila<br>Stock Center | BDSC:36599;RRID:BDSC_<br>36599 |
| *D. melanogaster*:<br>gnu RNAi | Kyoto Stock centre | HMC03185 |
| *D. melanogaster*: UAS-mRFP | Cytoplasmic RFP | Richa Rikhy, IISER Pune,<br>India |
| *D. melanogaster: Yp1-GFP* | Vienna stock centre | v318746 |
| *D. melanogaster: mTub>GFP-CAAX* | Plasma membrane marker, GFP tagged to CAAX | Yu-Chiun Wang as described earlier<sup>50</sup> |
| *D. melanogaster: UASp-Lifeact-TagRFP* | Bloomington Drosophila Stock Center | BDSC:58715;RRID:BDSC_58715 |
| *D. melanogaster: Utrrophin-mCherry* | sqh>Utrrophin-mCherry | Anna Sokac |
| *D. melanogaster: UASp-mEOS* | Bloomington Drosophila Stock Center | BDSC:604596;RRID:BDSC_604596 |

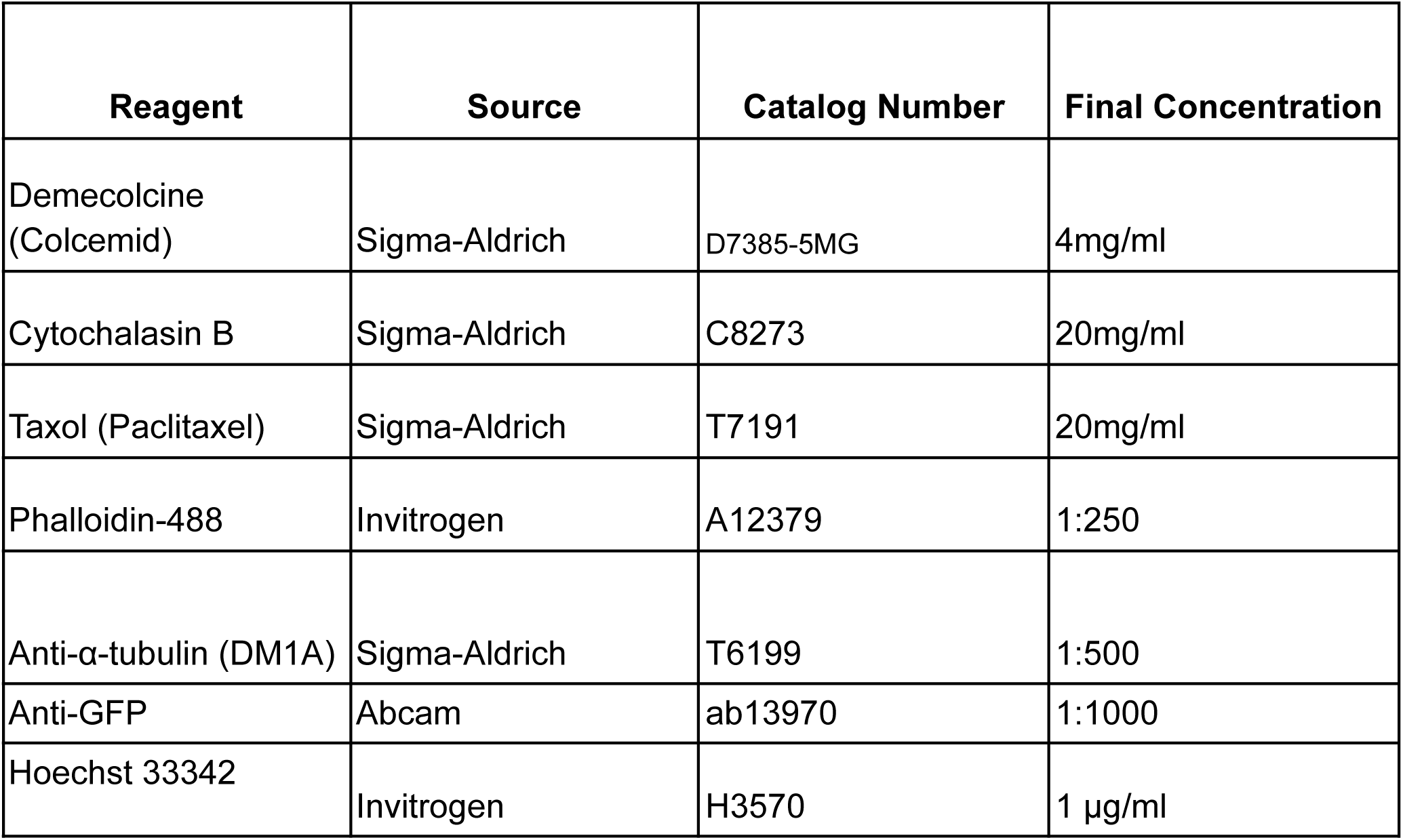
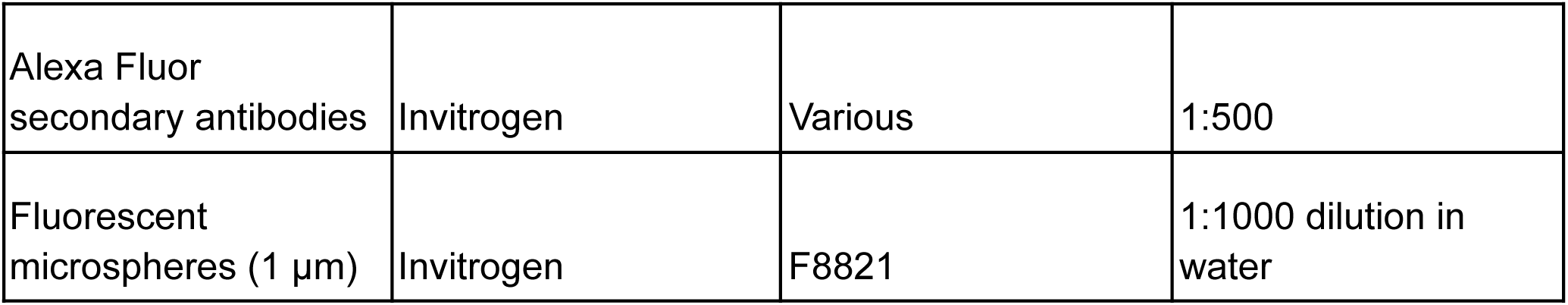

### Embryo Collection and Preparation

Embryos were collected from population cages maintained at 25°C using standard apple juice agar plates supplemented with live yeast paste. Collections were performed at 1.5-hour intervals to obtain embryos at appropriate developmental stages (NC 10-13).

For live imaging experiments, embryos were dechorionated using 4% sodium hypochlorite solution for 1 minute, followed by extensive washing with distilled water. Dechorionated embryos were mounted in 1× phosphate-buffered saline (PBS) for confocal microscopy.

### Live Confocal Microscopy

Live imaging of embryos was performed using two-photon scanning microscopy with a 25× water immersion objective (numerical aperture = 1.05) on an upright Olympus FVMPE-4GDRS system (InSight DeepSee pulsed IR Dual-Line laser, Spectra Physics) or an inverted Olympus FVRS-F2SJ system (Maitai and InSight DeepSee lasers). Excitation wavelengths were 930 nm for GFP and 1,040 nm (upright) or 1,100 nm (inverted) for mCherry, mRFP or mScarlet. Three imaging settings were used with the following parameters (*xy* dimension of the imaging region of interest (ROI), time interval, imaging angle or view): (1) ∼ 517 × 198 µm, 40 s, whole-embryo lateral views; (2) ∼ 209 × 81.80 µm, 2.75x zoom, 40 s, dorsal surface in the lateral views. For EB1 comet imaging, embryos expressing EB1-GFP and cytoplasmic mRFP were imaged on Nikon Ti2/Eclipse inverted microscope equipped and under a 60x/NA1.42 oil immersion objective using a Yokogawa CSU-W1/SORA imaging system. Two laser lines (488 and 561 nm) were used to excite the sample, while a tandem of sCMOS cameras (Prime BSI, Teledyne Photometrics) were used to acquire images with 2x2 binning.

### Drug Treatment Experiments

For injections, 0–1 hrs-old embryos were collected, dechorionated with bleach and mounted on an agar pad. The mounted embryos were then picked up using a coverslip painted with glue (prepared by immersing bits of Scotch tape in heptane), desiccated for 9 min using Drierite (W. A. Hammond Drierite Co.) and covered with a mixture of Halocarbon oil 700 and 27 (Sigma-Aldrich) at a ratio of 3:1. Needles for injection were prepared from micro-capillaries (Drummond Microcaps, outer diameter 0.97 mm, inner diameter 0.7 mm) pulled with a Sutter P-97/IVF and bevelled with a Narishige pipette beveller (EG-44). Injections were performed on a Zeiss Axio Observer D1 inverted microscope using a Narishige manipulator (MO-202U) and microinjector (IM300), followed by imaging. The following drug concentrations were used: Colcemid (4mg/ml in water), Taxol (20mg/ml in DMSO), Cytochalasin B (20mg/ml in DMSO). Control injections used equivalent volumes (∼65pL) of RNAse free water or DMSO as vehicle controls.

### Microsphere Injection and Tracking

To assess passive transport mechanisms, fluorescent polystyrene microspheres (1 µm diameter, Invitrogen) were injected into the cortical cytoplasm of living embryos. Microspheres were passivated as per a previous protocol ^51^, diluted 1:250 in water and injected using fine glass capillaries. Microsphere movement was tracked using time-lapse two-photon microscopy with 40-second intervals.

### Immunofluorescence Microscopy

For fixed sample analysis, embryos were collected at specific developmental stages and fixed in 4% paraformaldehyde for 20 minutes at room temperature. Following fixation, embryos were de-vittelinized using cold-methanol (for tubulin staining) or using hand-peeling (for phalloidin staining). Further staining methods and details were followed, as described previously ^24,52^.

### Image Analysis and Quantification

Image processing and analysis were performed using Fiji (ImageJ) ^53^ software with custom macros developed for this study. Further processing was carried out with custom made Python scripts.

#### Yolk segmentation

Yolk vesicle dimensions were quantified using a semi-automated image analysis pipeline. Z-stack images were first processed using a custom ImageJ macro that generated overlapping maximum intensity projections, reducing redundant detection of individual vesicles across multiple focal planes, followed by rolling ball background subtraction (radius 50) and Gaussian blur (σ = 2) to enhance vesicle contrast. These images were then segmented using either 1) automated threshold detection (Yen algorithm followed by default 16-bit thresholding), and the binary masks were refined through despeckle filtering, outlier removal, hole filling, and watershed segmentation to separate touching vesicles or 2) a self trained Cellpose model^54^. Individual vesicles meeting size criteria (1-70 μm² area) and circularity thresholds (0.8-1.0) were automatically identified and measured, excluding particles at image boundaries. Morphometric parameters including area, mean intensity, centroid position, and Feret diameter were extracted for each vesicle and saved as CSV files. Data from multiple embryos were pooled using a Python script.

#### Calculating the rate of ingression

To quantify yolk segregation dynamics (For example, Figure 1D), we generated kymographs from time-lapse images using a custom ImageJ macro. A 100-pixel line from the embryo cortex to the center was used with the Multi Kymograph plugin to create space-time plots. Kymographs were processed using Gaussian blur and intensity thresholding to generate binary images delineating the yolk-cytoplasm boundary. A custom Python script extracted the yolk front position at each time point, converted pixel coordinates to physical units, and cleaned the data by removing outliers, interpolating gaps, and applying smoothing. The processed distance-time data were exported, corrected for y-axis translations, and analyzed by simple linear regression in GraphPad Prism to determine segregation rates.

#### Measurement of the yolk depth from the embryo surface

To determine individual yolk vesicle positions relative to the embryo cortex (Figure 1G,4E,H, 5C,D,E), we manually drew and measured straight lines from the embryo surface to the edge of each vesicle using ImageJ. Multiple lines were drawn in an image, corresponding to each vesicle.

#### Calculation of yolk density

To quantify yolk vesicle density (Figure 1H), we used Yp1-GFP imaged embryos from Figure S1B, drew a box of 1024 X 180 (∼7700 μm^2^) starting at ∼200μm away from the cortex. Using TrackMate, GFP filled vesicles were identified as spots at each nuclear cycle. We computed the number of spots identified per 1000μm^2^ as yolk density.

#### Estimation of the yolk ingression speed from the segmented movies (MCE)

A detailed method is published in a companion paper in STAR Protocols. For the mass conservation equation (MCE), we start with the time-lapse movies obtained by applying the binary segmentation mask, showing the spatial variation of the yolk vesicles in the *xy* plane at four different *z* planes. Instead of the original fluorescence field, we now have an indicator function, which is represented by a binary occupancy field. If yolk is present, the pixel value is a constant non-zero value, else it is zero. We call it the yolk occupancy field ρ(*x*, *y*, *z*, *t*). Since, we are interested in obtaining the ingression speed along the *z* direction, we first integrate the occupancy field over the *xy* plane to obtain the yolk occupancy purely as a function of *z* and *t*, i.e ρ(*z*, *t*) = ∫∫ ρ(*x*, *y*, *z*, *t*)*dxdy*. The time evolution of ρ(*z*, *t*) can be expressed using the continuity equation:

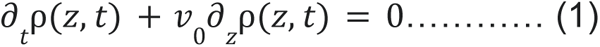

Here *v*_0_ is the ingression speed along the *z* −direction, which, for simplicity, we have assumed to be a constant. Furthermore, we have neglected any diffusional contributions to the currents. Equation 1 can be further integrated over the four *z* planes to obtain the total yolk occupancy number as a function of time which is defined as 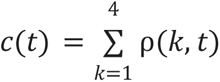. The time derivative of *c*(*t*) is then obtained from Equation (1) as

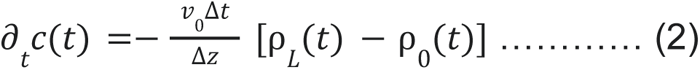

where ρ *_L_* (*t*) and ρ _0_(*t*) are the yolk occupancy fields at the bottom and top planes, respectively. Here Δ*z* is the distance between two consecutive *z* planes. Integrating Equation (2) further and normalizing with respect to the initial occupancy number *c*(0) with respect to time, we obtain an expression for the net yolk movement along the *z* direction, which is given as

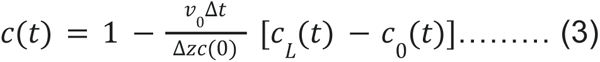

where 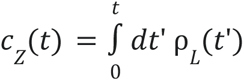 is the cumulative yolk occupancy number for a specific plane *z*, and *c*_0_ (*t*) and *c_L_* (*t*) denote the cumulative yolk occupancy number at the top (*z* = 0) and the bottom (*z* = *L*) planes respectively. In Equation (3), all the quantities in the RHS are known from experiments except for the ingression speed *v*_0_. We compare Equation (3) with the experimental *c*(*t*) profiles with *v*_0_ as the fit parameter. Note that this method gives a coarse-grained estimate of the ingression speed and does not account for local variations in the flow-field.

#### Quantification of cytoplasmic enrichment

mRFP fluorescence (Figure 1I, 4F) or photoconverted mEOS intensity (Figure 4G) was quantified from the time-lapse image series in Fiji/ImageJ. To prevent photobleaching of the signal, images were captured once every 10 minutes. On the mRFP channel, two rectangular regions of interest (400 × 50 px) were positioned over the cortex and over the yolk, together with a whole-image ROI, and the mean intensity of each region was measured in every frame ("Multi Measure"). The resulting per-frame intensity traces were exported and analysed in Python. To correct for global fluctuations (e.g. bleaching or acquisition changes), the cortical and yolk traces were normalized to the whole-image mean intensity at the same time point, giving a normalized intensity for each region. The intensity increase over the timelapse was then expressed as the normalized ratio of the last to the first time point (t_last/t_first) of each trace; a ratio of 1 indicates no change, and values above 1 indicate an enrichment.

## Statistical Analysis

All experiments were performed with biological replicates (n ≥ 3 embryos per condition) and technical replicates where appropriate. Data were plotted using GraphPad Prism 11.1.0. Statistical significance was assessed using appropriate tests given in the legends, with p < 0.05 considered significant. Data are presented as mean ± standard error of the mean (SEM) unless otherwise noted.

## Declaration of generative AI and AI-assisted technologies in the manuscript preparation process

During the preparation of this work the author(s) used Claude AI in order to polish grammar and sentence construction. After using this tool/service, the author(s) reviewed and edited the content as needed and took full responsibility for the content of the published article.

---

**Figure S1:**
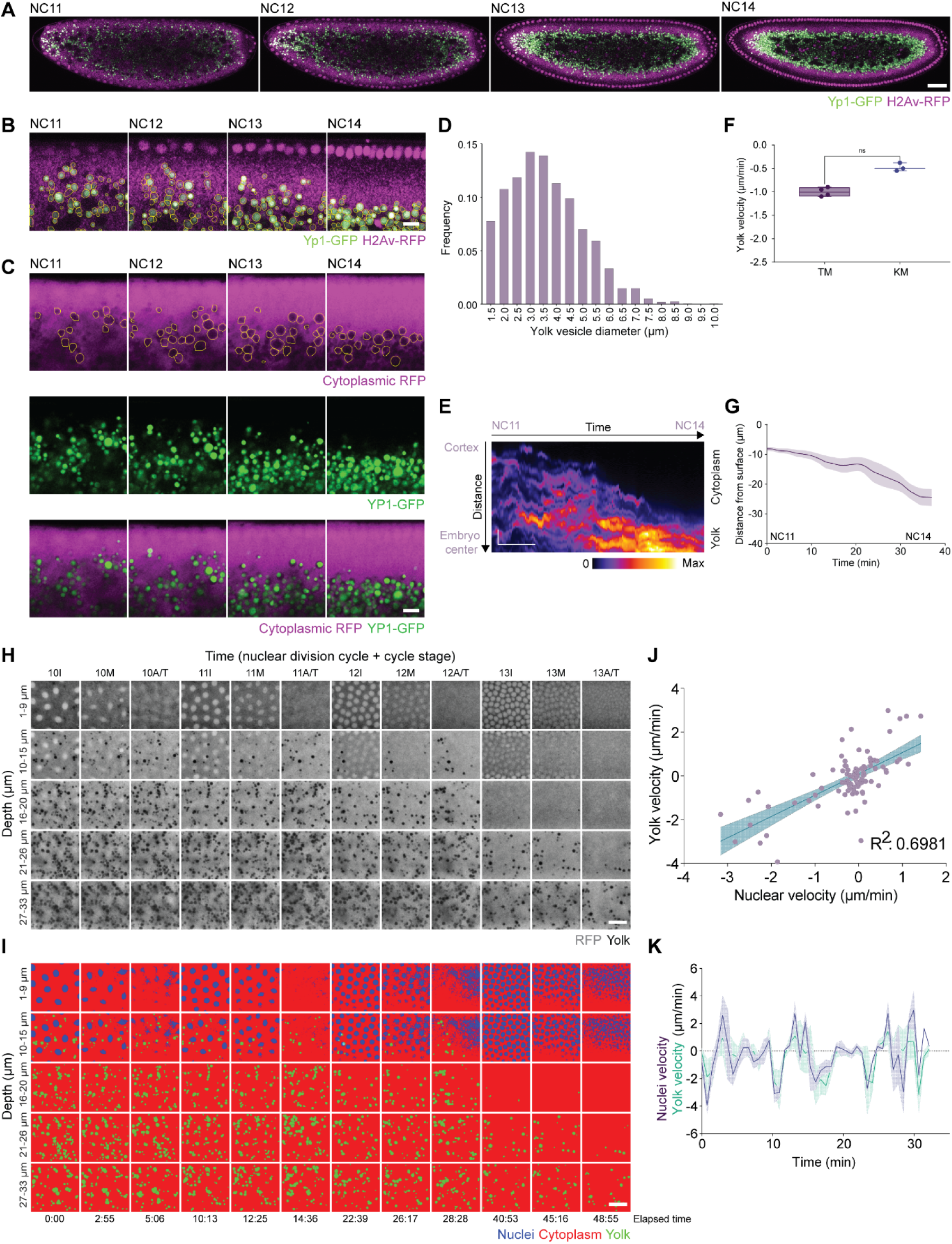
Quantitative analysis of yolk vesicle dynamics via YP1-GFP imaging. (A) Representative snapshots from live embryo imaging showing H2Av-RFP (which labels nuclei at the embryo cortex and lipid droplets in the embryo interior, magenta), Yp1-GFP filled vesicles (green), demonstrating the gradual inward movement of the yolk vesicles from the embryo surface toward the center from NC11 to NC14. Scale bar = 100 μm. (B) Higher magnification views of yolk vesicle dynamics from time-lapse imaging as in (A), with automated segmentation of the yolk vesicles outlined in yellow. Scale bar = 10 μm. (C) Snapshots of time lapse imaging with Yp1-GFP and cytoplasmic mRFP, revealing that Yp1-GFP labeled objects are identical to those negatively labeled with cytoplasmic RFP, thus validating the use of the negative labeling method. Scale bar = 10 μm. (D) Distribution of yolk vesicle diameters during pre-blastoderm stage, ranging from 1 to 9 μm, with a median of 3.43 μm (n=2073 yolk vesicles from 3 embryos). (E) A kymograph of Yp1-GFP derived from a cortex-to-center line profile, illustrating the temporal progression of yolk segregation from NC11 to NC14. Scale bar = 50 μm (vertical), 10 min (horizontal). (F) Quantification of the yolk velocities based on tracking of the yolk vesicles (“TM”, n=5 embryos) or linear fitting of the kymograph (“KM”, n=3 embryos). Mann-Whitney test. (G) Quantitative analysis of the yolk-cytoplasm boundary from NC11 to NC14 (n = 3 embryos, using segmentation and processing of kymographs as shown in (E). Error bars are mean±SEM. (H, I) Snapshots from XYZT imaging of an embryo expressing cytoplasmic RFP, displaying RFP distribution (H) and the corresponding segmented images (I) across a range of Z depths from NC10 to NC13 with cell cycle stages denoted as I, M, A/T for interphase, metaphase, and anaphase/telophase, respectively. Scale bar = 10 μm. Note that although this RFP construct does not contain any sequence motifs that could direct its subcellular localization, it has a slight bias towards the nucleus, but is fully excluded from the yolk vesicles. This allows for simultaneous visualization of nuclei, cytoplasm and yolk vesicles, following thresholding and segmentation, as shown in (I). (J) Nuclear velocity along an XY plane and yolk vesicle velocity measured from the nearest XY plane just underneath the nucleus showing a high degree of correlation (Pearson correlation coefficient, R^2^=0.69 (95%CI=0.57 to 0.79); n=3 embryos). (K) Nuclear velocity closely tracks that of the yolk vesicles in time, using velocity measured in the nuclear plane and the plane ∼10 μm below for yolk vesicle velocity, shown in (I).

**Figure S2:**
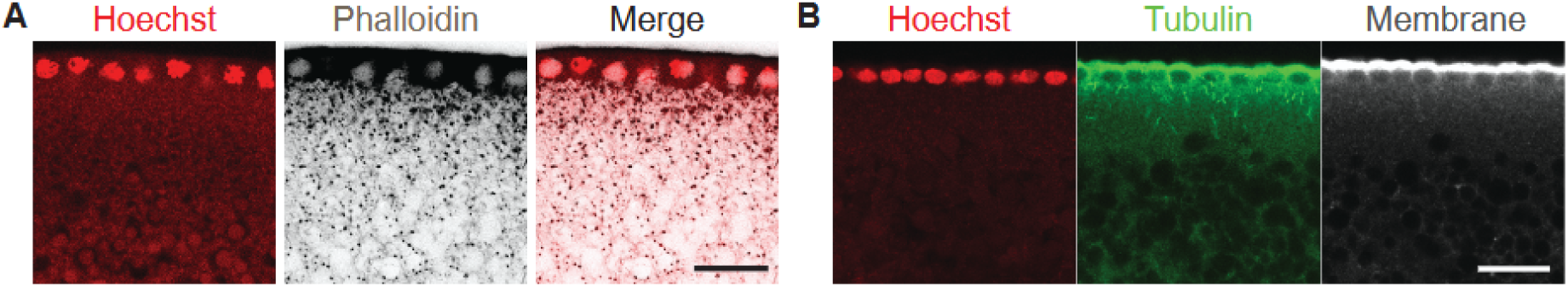
Yolk segregation and the concurrent cytoskeletal dynamics. Representative images of fixed embryos stained for (A) actin (phalloidin) and (B) tubulin (anti-tubulin antibodies). Yolk vesicles are negatively labeled in the phalloidin channel (A), or anti-tubulin and membrane channels (B). Oregon R embryos stained in (A), GFP-CAAX embryos in (B). Scale bar = 20 μm.

**Figure S3:**
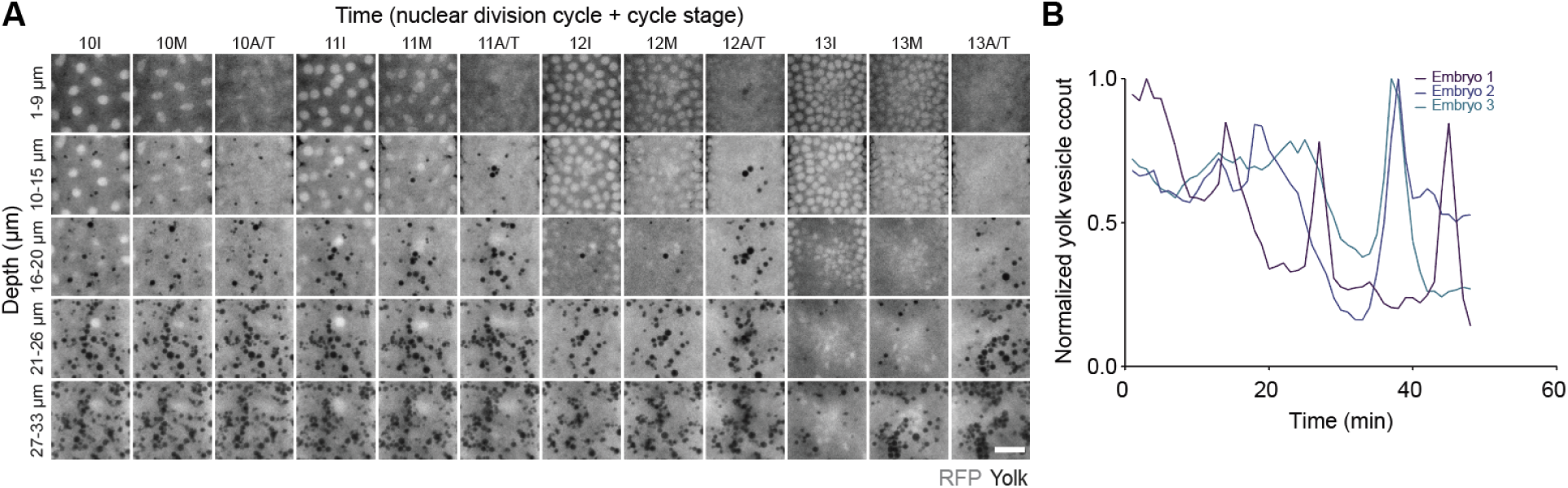
Yolk segregation is perturbed in the EB1 RNAi embryos. (A) Snapshots from XYZT imaging of yolk vesicles in an EB1-RNAi embryo labeled with cytoplasmic RFP, showing yolk distribution across Z depth from NC10 to NC13. I, M, A/T denotes cell cycle stages as interphase, metaphase, and anaphase/telophase, respectively. Scale bar = 10 μm. (B) Normalized yolk vesicle count as a function of time at the outermost 20 μm of embryo cortex in three representative EB1 RNAi embryos.

